# ESM-Scan - a tool to guide amino acid substitutions

**DOI:** 10.1101/2023.12.12.571273

**Authors:** Massimo G. Totaro, Uršula Vide, Regina Zausinger, Andreas Winkler, Gustav Oberdorfer

**Affiliations:** Institute of Biochemistry, Graz University of Technology, Petersgasse 12/2, 8010 Graz, Austria; BioTechMed Graz, Austria

## Abstract

Protein structure prediction and (re)design have gone through a revolution in the last three years. The tremendous progress in these fields has been almost exclusively driven by readily available machine-learning algorithms applied to protein folding and sequence design problems. Despite these advancements, predicting site-specific mutational effects on protein stability and function remains an unsolved problem. This is a persistent challenge mainly because the free energy of large systems is very difficult to compute with absolute accuracy and subtle changes to protein structures are also hard to capture with computational models. Here, we describe the implementation and use of ESM-Scan, which uses the ESM zero-shot predictor to scan entire protein sequences for preferential amino acid changes, thus enabling in-silico deep mutational scanning experiments. We benchmark ESM-Scan on its predictive capabilities for stability and functionality of sequence changes using three publicly available datasets and proceed by experimentally evaluating the tool’s performance on a challenging test case of a blue-light-activated diguanylate cyclase from Methylotenera species (*Ms*LadC). We used ESM-Scan to predict conservative sequence changes in a highly conserved region of this enzyme responsible for allosteric product inhibition. Our experimental results show that the ESM-zero shot model emerges as a robust method for inferring the impact of amino acid substitutions, especially when evolutionary and functional insights are intertwined. ESM-Scan is publicly available at https://huggingface.co/spaces/thaidaev/zsp

## INTRODUCTION

Proteins play a central role in various aspects of life, operating as fine-tuned molecular assemblies. The complexity of these structures is such that even a single alteration in the amino acid sequence can profoundly affect their functionality (Tokuriki and Tawfik 2009, Casadio et al. 2011, Zhang et al. 2018). Determining the amino acid substitution effects on protein structure, stability and function remains challenging despite continuous advances in structural and functional analyses (Shoichet et al. 1995, Fersht 1998, Gromiha 2007). Experimental approaches are often time-consuming and costly, even with high-throughput screening methods.

The earliest computational tools developed for this task leverage a combination of energy calculations, evolutionary insights and statistical analyses (Ng and Henikoff 2003, Yue, Li and Moult 2005, Smith and Kortemme 2011, Broom et al. 2017, Hopf et al. 2017). While achieving success rates in the modest range of 20% to 40%, these tools require substantial additional knowledge of the methods and the systems on the user’s part (Schmitz et al. 2020, Yamaguchi and Saito 2021, Hsu et al. 2022). Early machine learning approaches demonstrated good predictive power; however, they tended to overfit training data limiting their ability to generalise to proteins beyond the original training dataset (Bradford and Westhead 2004, Rao et al. 2019, Avery et al. 2022, Verkuil et al. 2022). Expanding beyond these early approaches, deep learning models harness vast sequence datasets and experimental information, autonomously learning from them in an unsupervised manner (Gilmer et al. 2017, Riesselman, Ingraham and Marks 2018). These models capture the intricate and non-linear contributions of individual components to protein stability and functionality (Ingraham et al. 2019, Sanderson et al. 2023). The generalisation capacity of deep learning models, however, remains an open topic, with persistent concerns about the risk of overfitting and the inconsistent transfer of learned knowledge to novel proteins beyond the training dataset (Notin 2022, Lin et al. 2023).

Arguably, the most well-known example of a deep learning model in biosciences is AlphaFold2, widely acknowledged as a transformative breakthrough in protein structural biology (Jumper et al. 2021, Tunyasuvunakool et al. 2021). Tweaks in the original model architecture endowed it with versatility, enabling it to undertake different tasks such as predicting multimeric assemblies, exploring alternative conformations and evaluating the impact of single amino acid substitutions on protein stability and function with varying levels of accuracy (Bryant, Pozzati and Elofsson 2022, McBride et al. 2022, Cheng et al. 2023, Sala, Hildebrand and Meiler 2023, Pak et al. 2023, Wayment-Steele et al. 2023). Another promising research area in deep learning explores language models. These models learn natural language syntax, semantics and contextual nuances and can generate new meaningful sentences. Drawing an analogy, protein sequences are akin to natural language texts, where amino acids correspond to letters, secondary structures to words, complete sequences to sentences and assemblies of several proteins to paragraphs (Ferruz, Schmidt and Höcker 2022, Lin et al. 2023, Wenzel 2023, Z. Zhang 2023). Protein-specific language models like ESM and BERT derivatives (Rives et al. 2021, Brandes, Ofer, et al. 2022, Chowdhury et al. 2022), based on the transformer architecture (Vaswani et al. 2017), are capable of learning and extrapolating hidden protein patterns that elude traditional energy-based methods (Chandra et al. 2023). An interesting aspect of these models is their potential use as zero-shot learners, predictors that can run on sets of classes not included in the training data without additional training (Larochelle, Erhan and Bengio 2008, Lampert, Nickisch and Harmeling 2009).

The outlined modelling approaches have the potential to transform day-to-day research for professionals working in biosciences. However, while some progress has been made, setting up and deploying an ML model is still non-trivial for most users. Moreover, data availability and curation are fundamental in the training and testing phases. The absence of a computational gold standard method currently hinders the direct transferability of model performances to varied use cases (Avery et al. 2022).

Here, we introduce ESM-Scan, a computational tool leveraging language models from the ESM family to infer rapidly and efficiently the fitness of amino acid substitutions on a given sequence (Meier et al. 2021, Brandes, Goldman, et al. 2022). Our tool is publicly available and requires no set-up from the user, ensuring ease of use. More skilled users can download the source code directly and adapt the tool for more specialised use cases. To illustrate the tool’s practical performance, expectations and limitations, we benchmark its performance against deep mutational scanning analysed by other computational tools described in the literature.

We further illustrate the tool’s efficacy through a practical application involving the blue-light-activated diguanylate cyclase from Methylotenera species (*Ms*LadC) as a well-characterised in-house model system (Vide et al. 2023). In this test case, we attempted to mitigate allosteric product inhibition of the GGDEF domain’s diguanylate cyclase activity (Schirmer 2016). This strategic approach aimed at improving our understanding of regulation mechanisms and enzyme functionality by increasing the abundance of active conformations. Preliminary experiments using amino acid substitutions suggested in the literature of related proteins (Teixeira, Holzschuh, Schirmer 2021) resulted in low expression yields and or non-functional proteins. This outcome is most likely due to the conserved region responsible for allosteric inhibition also being an integral part of an extensive inhibitory interface between sensor and effector domains that govern the inactive, dark state of *Ms*LadC (Vide et al. 2023). With ESM-Scan, we could identify promising alternative amino acid replacements suitable to generate protein variants potentially enriched in light state conformations for future detailed structural analyses of the activated state.

Overall, ESM-Scan offers a valuable and user-friendly resource to the scientific community, facilitating the prediction of amino acid substitution effects and empowering further specialised applications.

## METHODS

### ESM-Scan

Our ESM-Scan tool uses protein language models of the ESM family, trained as a masked language model (Devlin 2018). Operating as a masked language model means that each residue is assigned a probability score, based on its sequence context. Notably, a single residue alteration in an otherwise identical context produces a distinct score. Hence, fluctuations in these scores serve as indicators of the impact of amino acid substitutions. Positive scores signify enhanced fitness, whereas negative scores indicate the opposite. The latest iteration of ESM-Scan is hosted on HuggingFace (huggingface.co/spaces/thaidaev/zsp), featuring a user-friendly web interface that facilitates model inferences with minimal user configuration.

### Benchmarking datasets

The benchmarking datasets cover different use cases and originate from various experimental setups. For energy calculations, the Rosetta software suite stands as the industry standard (Das and Baker 2008, Kaufmann et al. 2010, Leaver-Fay et al. 2011). A score function, with statistics and physics-based calculations, approximates Gibbs free energy (ΔG). The ΔΔG, the distinction between the scores of a protein and any of its variants, serves as an estimate of the effects of the substitution, where negative values suggest structural stabilisation. ESM scoring is conducted using a local instance of ESM-Scan, in which the protein sequence and a list of amino acid substitutions are provided as input. The default parameters were used: the ‘esm2_t33_650M_UR50D’ model and the ‘masked-marginal’ scoring strategy.

**Dataset 1** derives from the work of Tsuboyama et al. (2023), which describes a high-throughput screening of protein variants accompanied by stability measurements. This dataset comprises data of more than 500 amino acid sequences, 32 to 72 residues long, folding as structural motifs or small domains. It includes both natural and Rosetta-designed *de novo* sequences from the Protein Data Bank (PDB). ΔΔG values are derived indirectly, by measuring resistance to trypsin/chymotrypsin. Our analysis used a total of 320000 unique data points from 420 distinct sequences in this dataset to compare measured vs. predicted ΔΔGs.

**Dataset 2** contains data on the expression levels and activity of Phosphatase and Tensin homolog (PTEN) variants. Cagiada et al. (2021) compiled this dataset to benchmark their coevolution-based metric against Rosetta-generated ΔΔGs and experimental findings from prior studies (Matreyek et al. 2018, Mighell, Evans-Dutson and O’Roak 2018, Suiter et al. 2020). PTEN data derived from studies evaluating cellular growth rate, drug sensitivity or cellular abundance. The first two metrics are classified as assessments of protein function, whereas the latter is an indicator of protein stability. The PTEN sequence, spanning 403 amino acids, underwent analysis with 5300 individual measurements on function and 7700 on structural integrity.

Furthermore, variant expressibility is binary-categorised as wild-type-like or non-wild-type-like. It is possible to identify threshold values for the ESM and ΔΔG scores that, when compared to this binary classification, accurately predict well-expressing vs lowly-expressing variants. This accuracy was measured by mapping Matthews correlation coefficient (φ) trends.

**Dataset 3** is a subset drawn from the SKEMPI and ZEMu databases (Moal and Fernández-Recio 2012, Dourado and Flores 2014), encompassing 66 multimeric proteins with nearly 900 recorded ΔΔGs for amino acid substitutions at protein-protein interfaces. Barlow et al. (2018) used it to develop energy-based methods for variant effect prediction. We calculated substitution ESM scores for individual chains and the multimeric complex. The complex assembly followed the procedure described by Lin et al. (2023), with 25-residue-long poly-G linkers separating chains, a protocol also employed in other studies (Mirdita et al. 2022, Tsaban et al. 2022).

Additionally, ESM scores were computed on a dataset frequently used for in-house testing in various approaches. Dehouck et al. (2009) curated this dataset to train a multi-layer perceptron for predicting ΔΔG using Rosetta-derived energy terms. Originating from the Protherm database (Gromiha 1999), it includes 2637 experimental measurements of single amino acid substitutions on 129 proteins.

The dataset analysis scripts are available at gitlab.tugraz.at/D5B8E35025578B91/esm-scan

### ESM-Scan application to *Ms*LadC

We ran ESM-Scan to perform a deep mutational scan of *Ms*LadC, whose sequence was obtained from UniProt entry A0A2S5LZS0 and adapted to the expression construct described in Vide et al. (2023). Utilising the ‘esm2b_t33_650M_UR50S’, identified as the optimal model for local execution on personal computers, we scored the variants of *Ms*LadC. We targeted Arginine 218 due to its role in allosteric inhibition and dark-state conformational stability.

### *Ms*LadC mutagenesis and protein production

Guided by ESM-Scan insights, we selected single-substitution *Ms*LadC variants R218K, R218S, R218N, R218A, R218V and R218D. These variants were generated by one-step site-directed mutagenesis, following the protocol described in Liu and Naismith (2008) using pET GB1a *Ms*LadC as a template (**Table S1** lists the primers used). The subsequent protein production and purification followed established protocols from Vide et al. (2023). Briefly, plasmids with the respective coding sequences were transformed into *Escherichia coli* BL21 (DE3) cells. Protein expression was induced with 0.1 mM isopropyl-β-D-thiogalactopyranoside at 16 °C for 16 hours after growth to mid-log phase in LB medium supplemented with kanamycin (0.03 g/liter). The harvested cells were lysed by pulsed sonication and the soluble fraction was affinity purified using Ni^2+^-Sepharose (Ni Sepharose 6 Fast Flow, GE Healthcare). The resulting *Ms*LadC variants, featuring N-terminal purification tags, were concentrated and subjected to size exclusion chromatography. Finally, the purified proteins were flash-frozen for storage at −80°C.

### *Ms*LadC functionality determination

For evaluating *Ms*LadC variant functionality, we assessed flavin cofactor binding and light state formation via UV-Vis spectroscopy and conducted an in vitro diguanylate cyclase assay to measure enzyme activity. Both analyses followed the established protocols outlined in Vide et al. (2023).

## RESULTS

Our results show a correlation between ESM scores and the observed impact of amino acid substitutions in proteins, demonstrating accuracies on par with state-of-the-art methods (Broom et al. 2020). Notably, the predictive capabilities of the model show an improvement when integrating protein functionality and evolutionary considerations into the scoring process. **Figure 1A** illustrates the prediction accuracies across the main datasets represented by the relative probability density function. In general, the more skewed each distribution is towards the top, the higher the linearity of the data.

**Figure 1:**
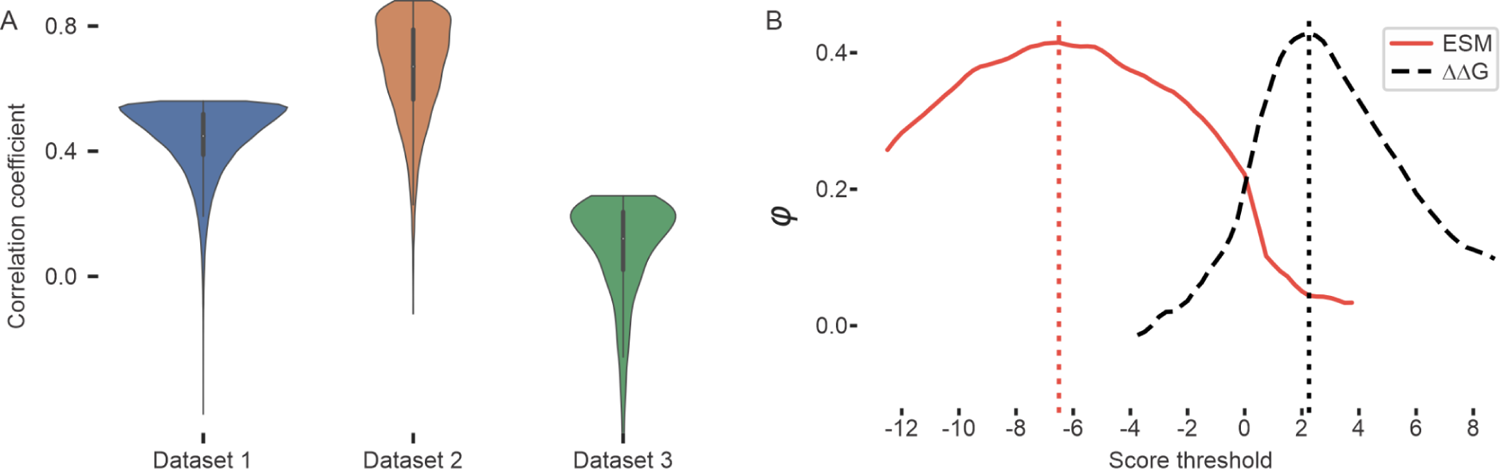
Benchmark correlations and functionality mapping. **A)** Correlation and error spread in benchmarked datasets. The distributions depict the probability density function of the normalised mean error for individual data points within each dataset. The means are adjusted to the calculated correlation R-value for enhanced visual representation. **B) Matthews correlation coefficient mappings for PTEN expression data.** Mapping of φ values at different ESM and ΔΔG score thresholds for PTEN data from dataset 2. The curve maxima correspond to the best score values to discriminate between well-expressing and non-well-expressing variants.

### Correlation between experimental, energy-based calculations and predicted **ΔΔ**G values

Dataset 1 shows an average correlation between the ESM scores and the energy differences measured due to substitutions. ΔΔGs are calculated indirectly through enzymatic digestion resistance, introducing some limitations in accuracy estimates. The overall correlation between ESM scores and ΔΔGs reveals a Pearson R coefficient of 0.44. This correlation value, albeit slightly low, is comparable to those obtained by traditional methods for estimating ΔΔG. A direct comparison on the same dataset using Rosetta calculated ΔΔG values is impractical due to its computational demand, highlighting a notable advantage in favour of our tool.

Dataset 2 offers functional and protein expression level measurements for PTEN to compare with Rosetta-calculated ΔΔGs and coevolutionary scores. ESM scores exhibit a correlation with protein abundance (R = 0.48), demonstrating a performance that is comparable to Rosetta ΔΔGs (R = 0.49) and coevolutionary score (R = 0.48). When considering PTEN functionality, there is a marginal decrease in prediction accuracy for Rosetta and coevolutionary scores (R = 0.42 and R = 0.46, respectively), while ESM outperforms significantly with an R-value of 0.56.

Scanning the ESM and ΔΔG score threshold domains and correlating them with the binary classification for PTEN expression, we could map trends in φ values, as depicted in **Figure 1B**. The peaks in the curve correspond to threshold values that optimally discriminate between wild-type-like-expressing and lowly-expressing variants. For instance, an ESM score of −6.5 emerges as the most accurate discriminator. Notably, these findings align with energy-based reference ΔΔG values, affirming the destabilising effects of amino acid substitutions in the range of 2-3 kcal/mol.

In contrast to the R-value of 0.58 reported by Barlow et al. (2018) for their Rosetta-based analysis system, ESM-Scan demonstrates a notable lack of correlation with experimental results for protein-protein interfaces in dataset 3. Even considering the substitutions in the context of multimeric assemblies, the predictive power of ESM does not improve. In fact, the effect on correlation coefficients decreasing from 0.09 to 0.06 is likely statistically insignificant. It’s crucial to note that the model used in this study differs in architecture and size from the ESMFold model described by Lin et al. (203) despite belonging to the same family.

The correlation between ESM scores and the heterogeneous data from Dehouck et al. (2009) reveals only a modest R-value of 0.17. This result, while lower than expected, probably relates to the fact that a variety of protein origins and experimental settings are combined in the dataset. Considering the good correlation with the datasets mentioned above, one should be cautious in overinterpreting the prediction misalignment in this specific case.

### Test case *Ms*LadC

Based on the promising results obtained in the benchmarking datasets, we applied ESM-Scan to an in-house model system, *Ms*LadC (**Figure 2A-C**). We aimed to generate protein variants that retain wild-type functional properties but lack allosteric product inhibition. We focused on Arginine 218 (**Figure 2A,B**), located within the characteristic inhibitory site (aa-sequence motif RxxD), which is intricately involved in the coordination of c-di-GMP dimers as part of the diguanylate cyclases’ feedback inhibition mechanism (Christen et al. 2006, Schirmer 2016). Leveraging insights from the ESM-Scan, we chose to express the *Ms*LadC R218K, R218S, R218N, R218A, R218V, and R218D variants. Despite relatively consistent total expression levels across these variants (**Figure S1A**), solubility and functionality were significantly affected (**Figure S1B** and **Table S2**), highlighting the importance of Arginine 218 in *Ms*LadC and confirming the tool’s negative scores for this position (**Figure 2B,C** and **Figure S2**). ESM rankings, purification success, and protein functionality correlate well, with all variants consistently underperforming compared to the wild-type (**Table S2**).

**Figure 2:**
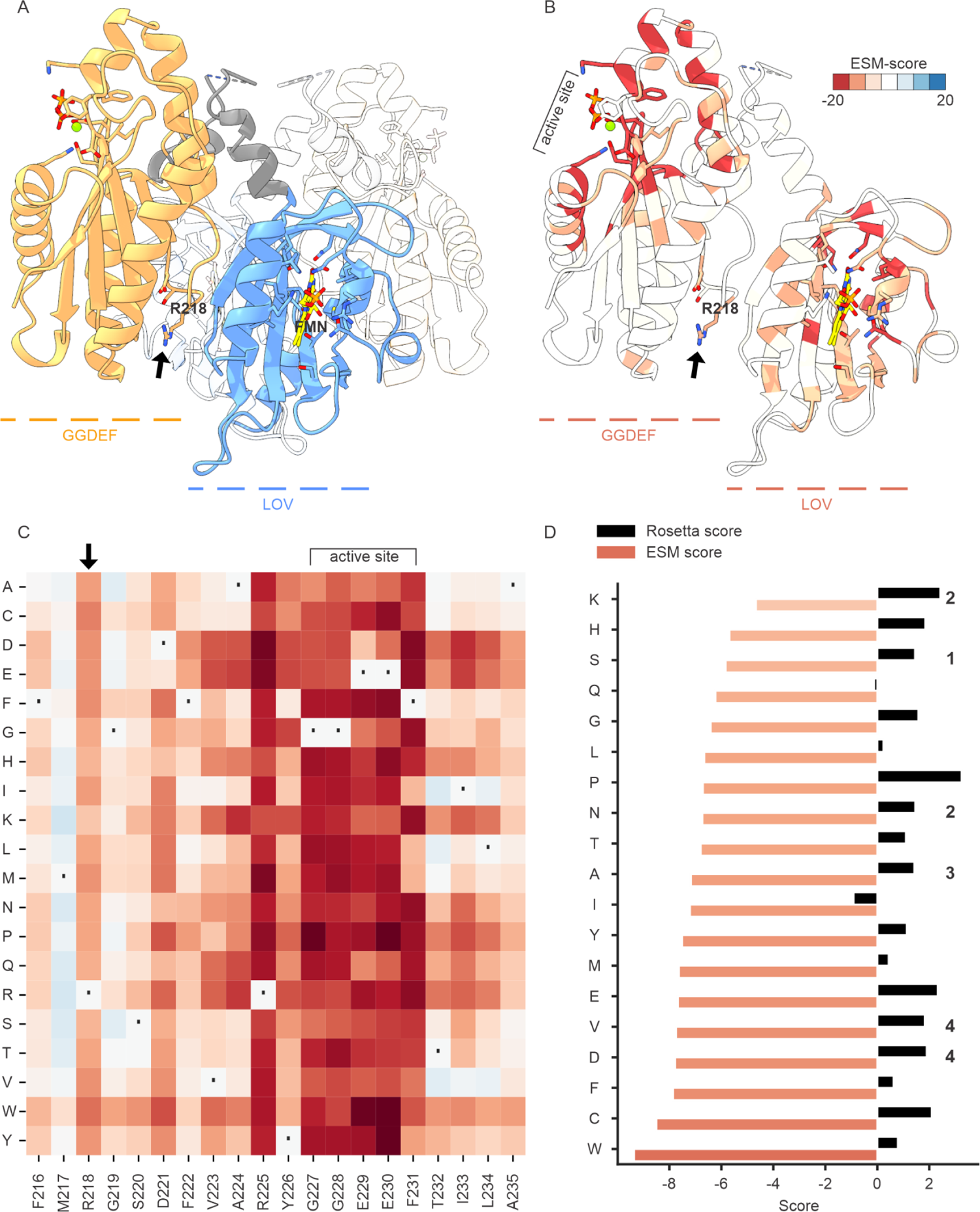
Test case MsLadC. **A,B**) Cartoon representation of the MsLadC crystal structure in its dark, inactive state (PDB 8C05). The residues R(218)xxD, lining the inhibitory interface between the LOV and GGDEF domain, are shown as stick models and R218 is indicated by an arrow. The FMN cofactor (yellow), its binding site, and the active site with pyrophosphate (orange) and a Mg^2+^ ion (green) are shown as stick models. In **A)**, the functionally relevant dimeric state is shown with one LOV domain in blue, the linker helix in grey, the GGDEF domain in orange, and one protomer distinguished in transparency. **B)** shows a single protomer coloured according to the ESM-Scan scores heatmap, panel C, averaged at each residue position and binned into seven bins. **C**) The ESM-predicted fitness landscape for the MsLadC deep mutational scan at positions 216-235. The unbinned, continuous colour gradient matches the range defined in panel B. The complete heatmap for all residues is shown in **Figure S2**. A dot indicates wild-type residues and an arrow points to R218. **D**) Comparison of ESM and Rosetta scores for R218 substitutions. Higher ESM values indicate fitter variants, whereas the opposite is true for Rosetta scores. The number next to the columns indicates the experimentally tested variant ranking based on **Table S2**.

Post-purification, protein yields for the top-ranking R218K, R218S, and R218N variants were five times lower than the wild-type, diminishing to 15 times lower yield for R218A. R218V and R218D even lost the ability to bind the FMN cofactor of the blue-light sensing LOV domain of *Ms*LadC, resulting in sensory function loss. In the case of the latter two variants, impairment in sensory function was already visible during cell harvest, as the pellets lacked the typical yellow colour. While the protein of interest could still be detected on SDS-PAGE of purified soluble fractions (**Figure S1B**), the low levels precluded any enzymatic activity assessment. Enzymatic assays for other variants indicated significantly reduced diguanylate cyclase activity compared to the wild-type (**Table S2**). A comparison with the state-of-the-art Rosetta-calculated energies for potential substitutions at position 218 reveals a lack of correlation with the experimental data (**Figure 2D**).

Our empirical observations from the *Ms*LadC test case align with the observed ESM thresholds calculated for dataset 2 (**Figure 1B**), where values below −6.5 indicate non-viable substitutions (**Figure 2D**).

## CONCLUSIONS

Estimating the effect of point mutations on protein stability and function is still challenging. Here, we introduce ESM-Scan, a user-friendly tool designed to predict amino acid sequence variants. Our analysis, comparing predictions to experimental data from deep mutational scanning experiments, offers valuable insights into the potential and limitations of this tool.

ESM-1v, the original model developed by Meier et al. (2021), demonstrates a certain ‘understanding’ of protein grammar, reporting correlations to experimental results of around 0.5, a value that stands favourably against state-of-the-art models. Our findings point to similar performances when predicting effects on the stability of monomeric proteins from large datasets, on average. However, ESM-Scan can yield low-accuracy predictions, in particular, if the data include significant differences in experimental setup or ΔΔG calculation. Variable performance should be anticipated, as depicted in **Figure S3**. Moreover, even though ESM models have been used to model multimeric complexes, predictions at protein-protein interfaces appear unreliable. Whether this is due to model discrepancies, unfitness or dataset biases, we cannot make a case for this particular use and we encourage considering alternative tools (X. Liu et al. 2021, Cagiada et al. 2023, Sieg and Rarey 2023).

ESM shows consistently better performances when protein functionality is part of the equation. Its inferred values align closely with those provided by co-evolutionary analysis tools and, in some instances, even surpass them. This ability might be confined to naturally occurring proteins or domains included during training; future *de novo* protein functionality assays might allow testing this hypothesis.

Deep learning tools trained on structural information are likely superior predictors in tasks where spatial features are pre-eminent, as in the case of protein-protein interactions (Wang et al. 2022, Heinzinger et al. 2023, Shu et al. 2023, Su et al. 2023). AlphaFold2 and ProteinMPNN, a model developed around a message-passing neural network, have shown potential in predicting fitness variations upon residue substitution (Dauparas et al. 2022, Brown et al. 2023, Reeves and Kalyaanamoorthy 2023). Similar architectures can model conformational free energy surfaces with reasonable approximation, although challenges persist in efficient structure representation and learning (Jing et al. 2020, Ovchinnikov and Huang 2021).

In our test case, ESM-Scan proved instrumental in guiding *Ms*LadC substitution variants of a critical amino acid involved in two different functionalities - product inhibition and inactive dark-state assembly formation. Our initial literature-guided rational approach failed to yield viable protein variants for thorough *in vitro* characterisation due to a substantial impact on protein yield. ESM-Scan helped us generate functional *Ms*LadC variants and ESM scores correlated strongly with the experimental findings regarding protein solubility and functionality. Furthermore, a retrospective examination of previously published data on *Ms*LadC variants (Vide et al. 2023) reveals a consistent trend: ESM-Scan scores negatively for non-viable variants. Negative scores generally cluster around functional residues and flanking regions (**Figure 2B,C**); however, choosing the best-ranked substitution is likely to outperform the alternatives. While this opens opportunities for further investigating R(218)xxD-motif-mediated allosteric product inhibition, such efforts go beyond the scope of this work.

Integrating the ESM model’s zero-shot predictor into an accessible interface generated ESM-Scan, an *in-silico* deep mutational scanning tool. Geared for the casual user, it operates online without additional setup, computational resource management, or coding expertise. With swift inference times and minimal overhead, the tool is well-suited for preliminary screenings. Our benchmarking on independent datasets aligns with expected performances. Moreover, we demonstrate the application of ESM-Scan for conducting an *in-silico* deep mutational scan to modulate the functionality of *Ms*LadC. As in any *in-silico* prediction, we want to emphasise that the user must be mindful when interpreting results, testing the model behaviour in each case and adjusting threshold values. For adept users, the flexibility to adapt ESM-Scan for self-hosted systems offers possibilities for fine-tuning to individual needs and potentially superior results in specific instances.

In essence, the ESM model emerges as a robust method for inferring the impact of amino acid substitutions, especially when evolutionary and functional insights are intertwined. Its inferential capabilities present an exciting avenue for *in-silico* prediction of protein functionality alterations, potentially reducing the need for resource-intensive wet lab experiments.

## ACKNOWLEDGEMENTS

MT and UV are supported by the Austrian Science Fund (FWF) grant 10.55776/DOC130 and MT additionally by the Styrian Government (Zukunftsfonds, doc.fund program). MT and UV were trained within the framework of the PhD program Biomolecular Structures and Interactions (BioMolStruct). GO was supported by funding from the European Research Council through a Starting Grant (HelixMold 802217). This research was funded in whole, or in part, by the Austrian Science Fund (FWF) [10.55776/P34387 to AW and 10.55776/P30826 to GO]. For open access, the authors have applied a CC BY public copyright licence to any Author Accepted Manuscript version arising from this submission.

## SUPPLEMENTARY MATERIALS

**Figure S1:**
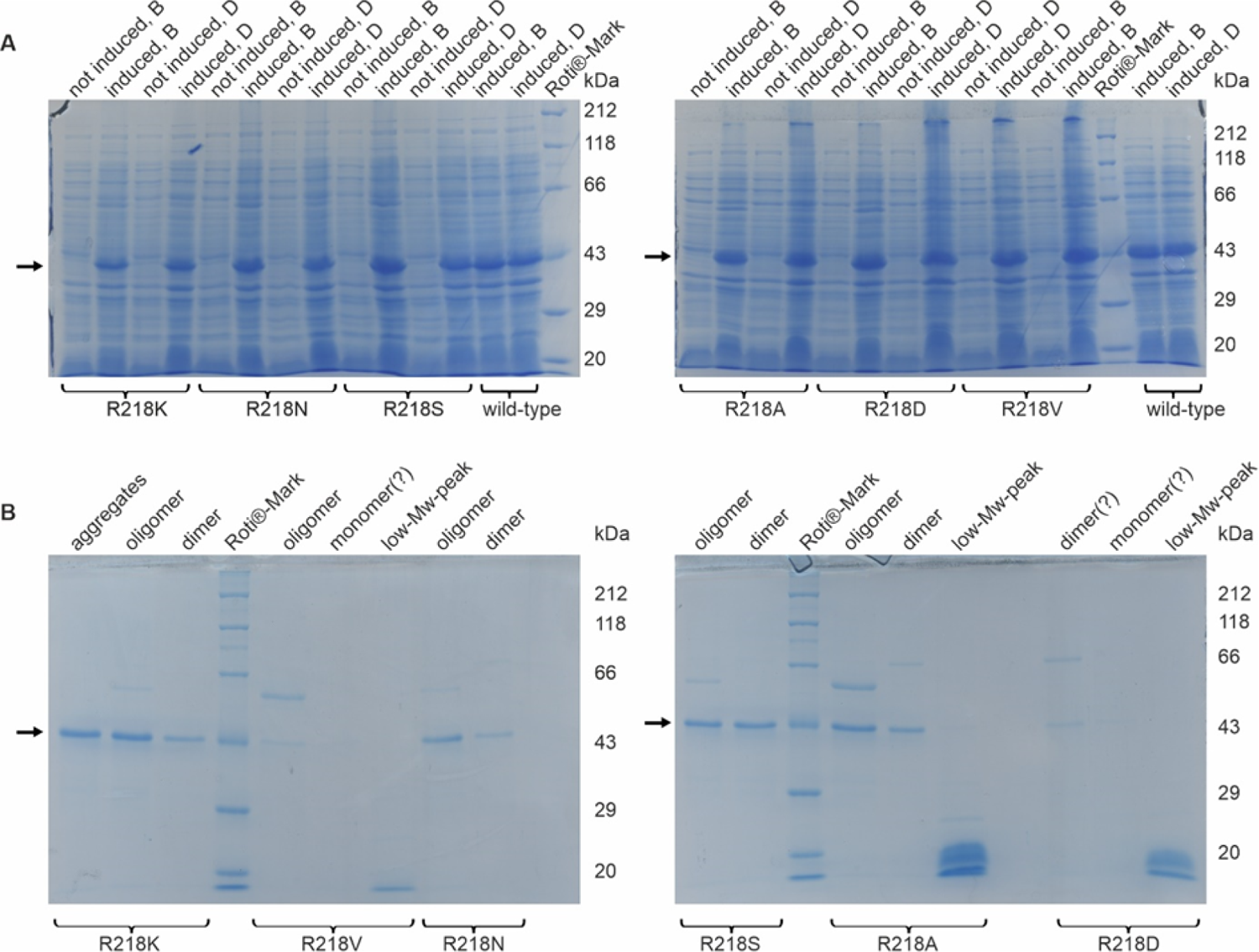
Analysis of *Ms*LadC-R218 variants overexpression and purification yield. SDS-PAGE (12.5%) gels in panel **A**) show samples before and after induction with IPTG, both under continuous blue-light illumination (B) and under dark conditions (D). In panel **B**) size-exclusion fractions were analysed, corresponding to aggregates, oligomers, dimers, monomers or other low-molecular-weight molecules.

**Figure S2:**
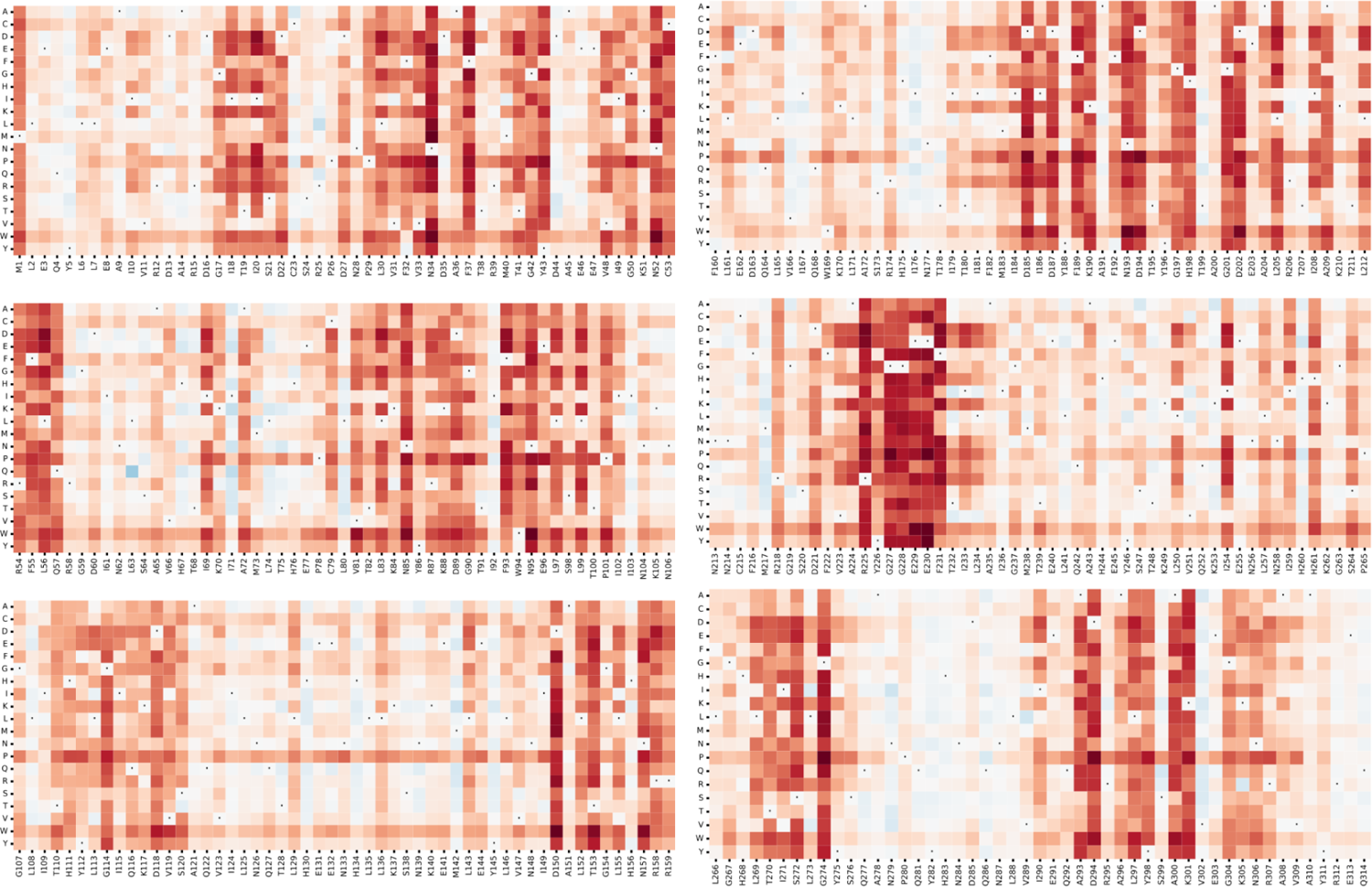
Deep mutational scanning of *Ms*LadC. For every residue position of *Ms*LadC, all possible alternative amino acids are scored with the default ESM-Scan parameters. The scores are colour-coded in blue for positive values and red for negative ones, in the range of −20 to 20. The wild-type amino acid scores at zero and is marked by a dot.

**Figure S3:**
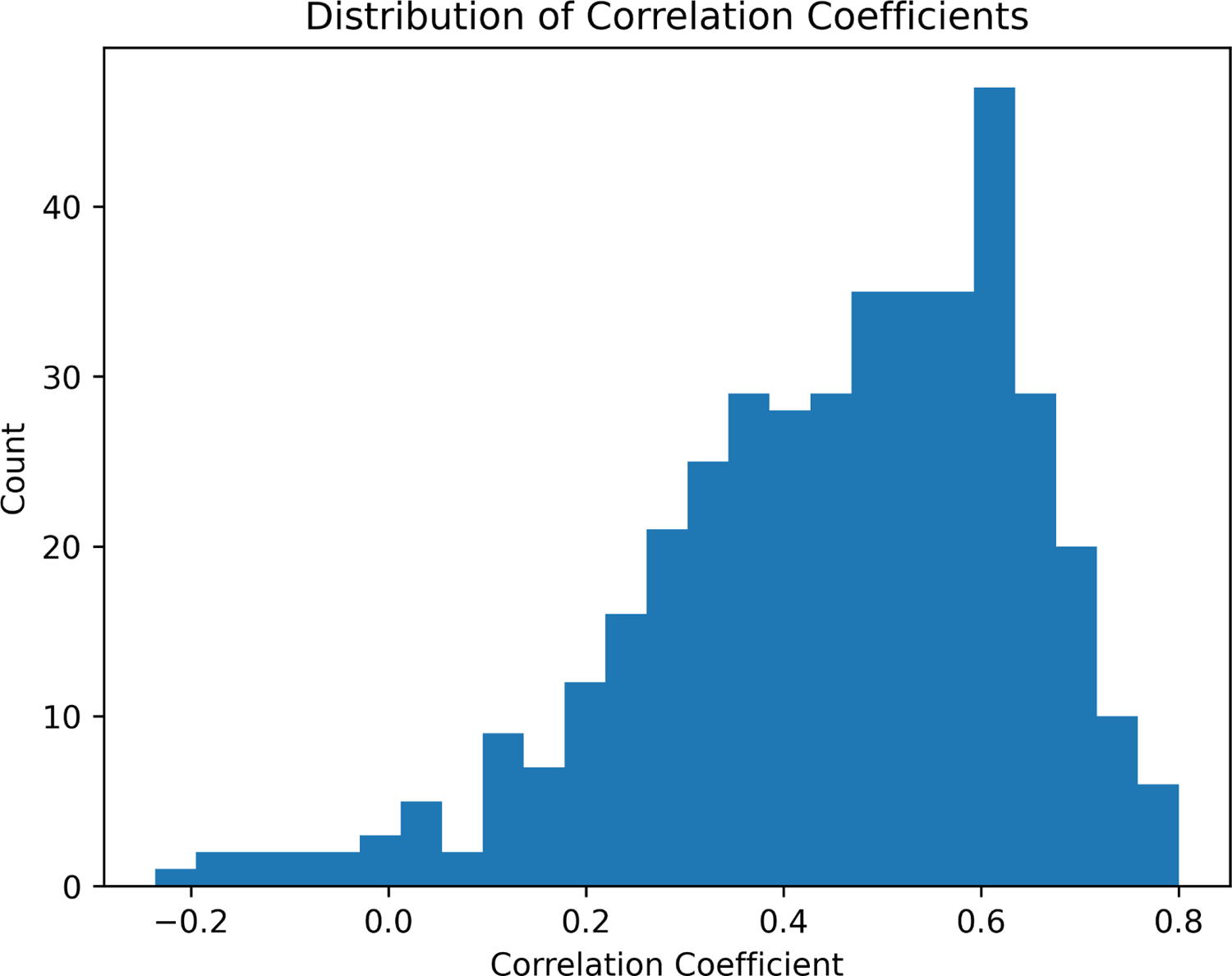
Distribution of correlations in Dataset 1. The graph shows the distributions of correlations of all unique sequences in dataset 1. For each of the x sequences the deep mutational scanning yields a correlation value plotted on the x-axis; the sets are unweighted, contributing to one unit on the y-axis independently of the number of experimental data in each.

**Table S1:**
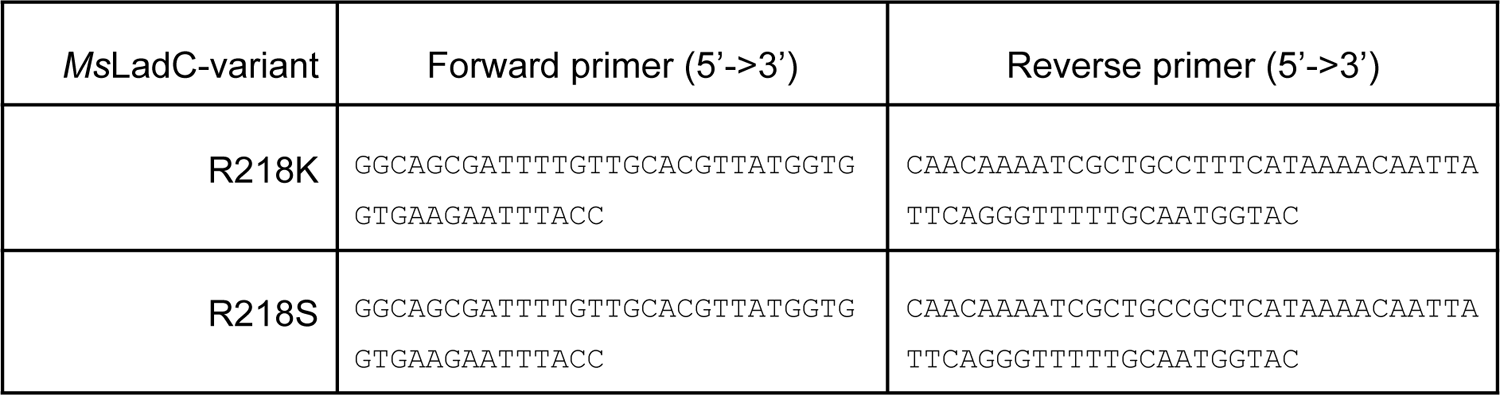
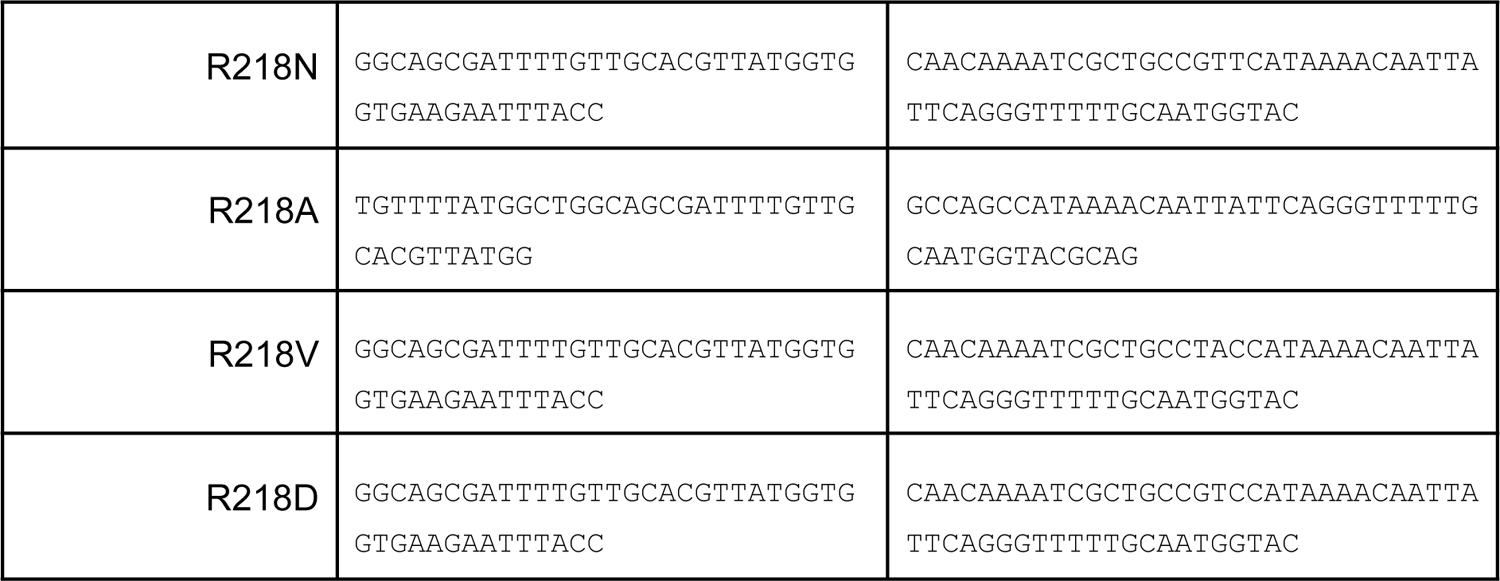
Primers for site-directed mutagenesis of *Ms*LadC.

**Table S2:** *Ms*LadC-R218 variants’ overexpression and purification efficiency. This legend categorises expression efficiency, FMN-bound protein yield and enzymatic activity, determined from the initial velocities of the light-state diguanylate cyclase activities, compared to wild-type behaviour. Symbol Key: ✓ wild-type behaviour; - no functional protein yield.

| *MsLadC*-variant | Overexpression | Yield compared to wild-type<br>(based on FMN binding) | Diguanylate cyclase<br>functionality compared to wild-type |
| --- | --- | --- | --- |
| R218K | ✓ | 0.2x | 0.1x |
| R218S | ✓ | 0.2x | 0.5x |
| R218N | ✓ | 0.2x | 0.2x |
| R218A | ✓ | 0.07x | 0.01x |
| R218V | ✓ | - | not measured |
| R218D | ✓ | - | not measured |

